# Deep Learning for Multi-task Plant Phenotyping

**DOI:** 10.1101/204552

**Authors:** Michael P. Pound, Jonathan A. Atkinson, Darren M. Wells, Tony P. Pridmore, Andrew P. French

## Abstract

Plant phenotyping has continued to pose a challenge to computer vision for many years. There is a particular demand to accurately quantify images of crops, and the natural variability and structure of these plants presents unique difficulties. Recently, machine learning approaches have shown impressive results in many areas of computer vision, but these rely on large datasets that are at present not available for crops. We present a new dataset, called ACID, that provides hundreds of accurately annotated images of wheat spikes and spikelets, along with image level class annotation. We then present a deep learning approach capable of accurately localising wheat spikes and spikelets, despite the varied nature of this dataset. As well as locating features, our network offers near perfect counting accuracy for spikes (95.91%) and spikelets (99.66%). We also extend the network to perform simultaneous classification of images, demonstrating the power of multi-task deep architectures for plant phenotyping. We hope that our dataset will be useful to researchers in continued improvement of plant and crop phenotyping. With this in mind, alongside the dataset we will make all code and trained models available online.

## 1. Introduction

Crop phenotyping performs a crucial role in the development of higher-yielding plants, which in itself offers one solution to the continuing challenge of global food security. For cereal plants, yield is measured in terms of grain, found within the spikes at the tip of the plant. Therefore, counting both the number of spikes, and the so-called spikelets within them (see Fig. 8) is an important measure. In this work, we aim to further the state-of-the-art in wheat phenotyping. We present a dataset, publicly available, that can be used by researchers to improve their wheat phenotyping ability via machine learning. Using this dataset, we apply a deep network architecture to perform simultaneous localization of the spikes and spikelets, and image classification. The result is a system capable of counting and locating both spikes and spikelets, in the presence of varied phenotypes, occlusion, clutter, and arbitrary rotations. Example outputs of our approach can be seen in Fig. 1. The dataset and all related code can be found at *http://plantimages.nottingham.ac.uk/*.

**Figure 1:**
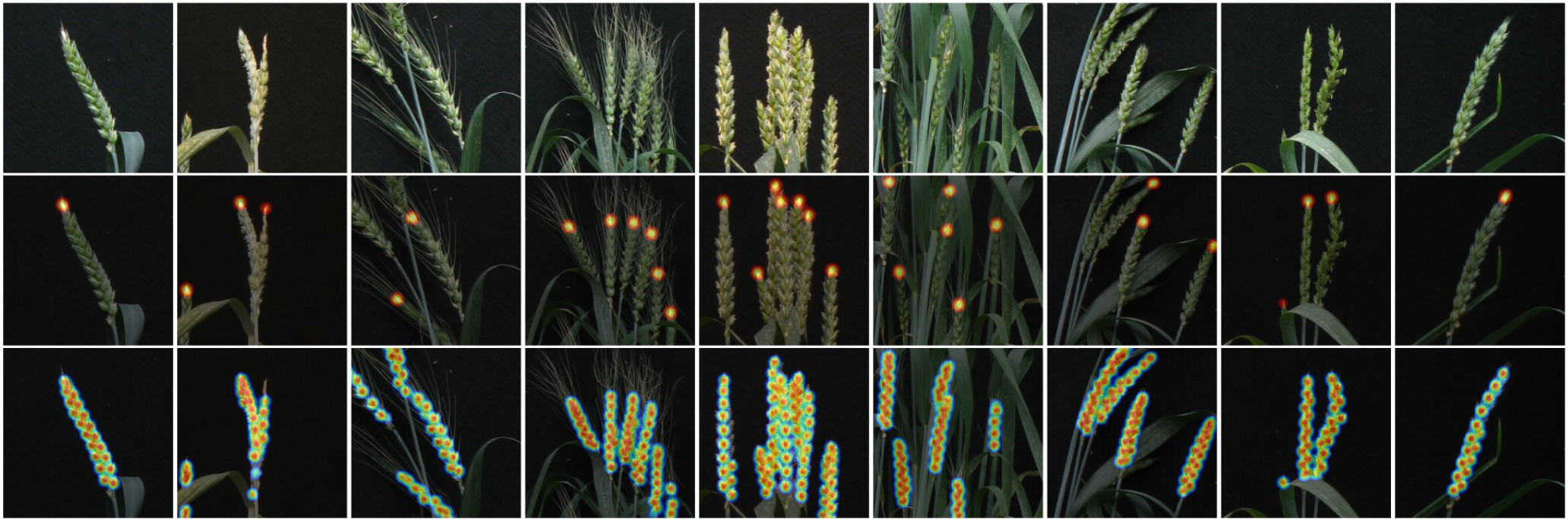
A selection of results from our deep network locating spikes (middle) and spikelets (bottom) on the ACID dataset

**Motivation.** Biologically, the ability to phenotype spike traits is of great importance, with uses in several different research areas. For example, spikelet development and thus yield can be affected by abiotic stresses such as high temperature and drought [12, 26] or changes in sowing date [1]. Other traits such as the presence of awns, a bristle or hair like structure extending from the end of each floret (see Fig. 2), have been linked with increased photosynthesis under drought and greater harvestable yield [4, 19]. Intricacies within the spike and features of the spikelets themselves have been shown to be useful when predicting yield [14].

**Figure 2:**
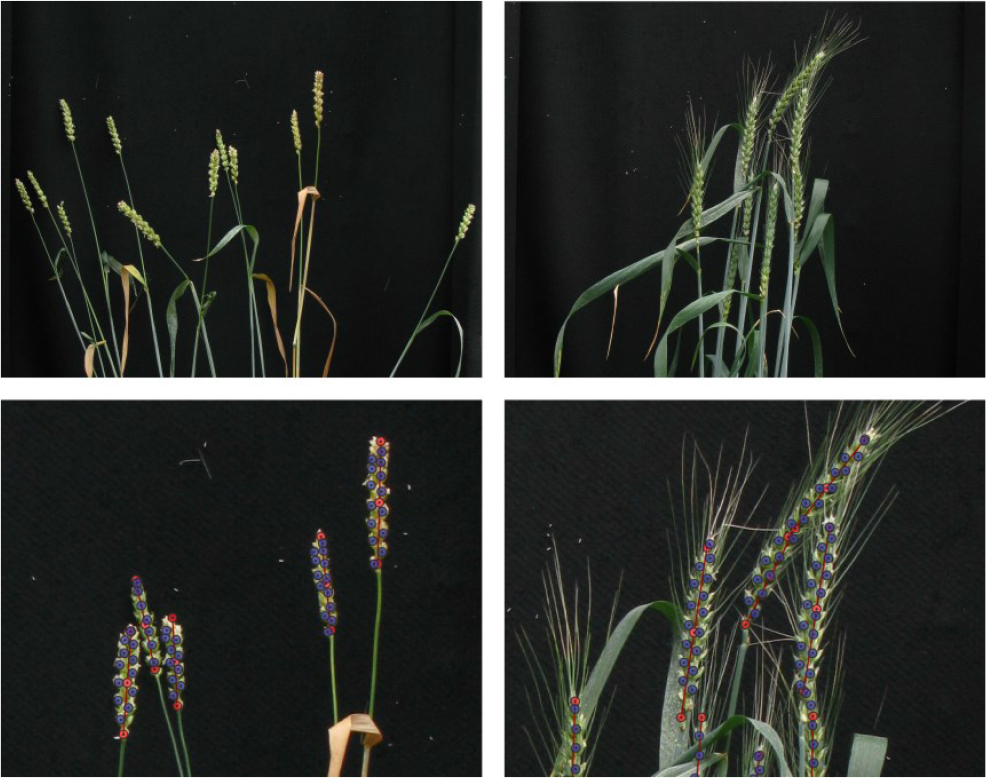
Representative images from the ACID dataset. Note presence of awns (bristles) in the right-hand images. Expert annotations are shown.

Locating and counting spikes, and particularly the spikelets within, presents a demanding computer vision challenge, as can be seen in Figs. 1, 2 and 9. Occlusion, self similarity, variation in appearance, viewpoint sensitivity and lighting challenges weigh heavy on the ability of traditional image analysis to satisfactorily solve this task. But these challenges are not unique to wheat spikes; rather they are representative of general plant and crop phenotyping challenges that present themselves when we start to move from the lab to the field. Therefore, by addressing this specific, challenging task, we are harnessing deep learning to solve a realistic phenotyping problem which if successful bodes well for other phenotyping automation felt beyond the reach of traditional analysis. This then is our acid test of deep learning in the wild.

### 1.1. Contributions

We present a new dataset (which we call ACID: the Annotated Crop Image Dataset) of accurately labeled wheat images, and adapt an appropriate CNN architecture to address the challenge of wheat spike phenotyping. Our network performs multi-task learning, simultaneously locating spike features, whilst also classifying the awned phenotype of the wheat in each case. An overview of our network architecture is shown in Fig. 3. In summary, our contributions are:

- A new publically-available dataset containing 520 images of wheat plants exhibiting a wide range of canopy and spike phenotypes. Each plant has been carefully annotated by an expert, including the locations of each spike, and individual spikelets. Each image has also been classified as exhibiting, or not, an awned phenotype.
- A novel application of a deep CNN architecture for multi-task learning in wheat phenotyping. We have extended a state-of-the-art CNN architecture to perform simultaneous localisation of features and classification of images. This network is trained end-to-end from scratch on the ACID dataset, and we present detailed discussions of our training process, along with results on its performance.
- An adapted data augmentation approach to handle views of multiple similar objects. Unlike many existing image sets, ACID contains many instances of the same class of object, with extremely similar appearance. This presents a unique challenge to the traditional network training and data augmentation approaches seen previously. We present a spike-centric data augmentation approach that varies the data between epochs as much as possible, leading to improved results.

## 2. Related Work

Advances in image-based plant phenotyping helped drive plant research for many years. Traditionally, low-level image processing approaches were the norm, using pixel-based techniques and hand crafted models to aid localization and measurement of plants (for example [8, 2]). Recently machine-learning based approaches have seen increased adoption, allowing systems to learn how to find and segment plants, based often on a hand-crafted feature set (for a thorough overview of machine learning approaches in plant phenotyping see [22]). Most recently, deep learning promises a step-change in the performance of many image-based systems (e.g. [18], [23]), but adoption by the plant phenotyping community is still in its infancy.

To date, most spikelet and ear counting is done by hand, following a method similar to [14], which relates crop yield to features in the spike and spikelets without using image analysis. Some methods do exist for automatically detecting heading and flowering in wheat - [21] uses a bag-of-visual-words approach to identify growth stages in field-grown wheat. Low level features are extracted using the SIFT algorithm. Finally support vector machine classification is used to classify growth stage. Accuracies for stage classification range from 85% (flowering) to 99% (late growth stage). Colour and texture have been used in a preliminary study to count wheat ears [7], reporting accuracy up to 85%, although, as the authors concede, this is across a small and limited dataset, and relies on a bespoke image processing pipeline.

Machine learning approaches have been applied recently to a number of other plant phenotyping challenges. Some cereal-specific examples include using expectation maximization to identify wheat streak mosaic virus[6]; Simplex Volume Maximisation to discover characteristic spectra in hyperspectral data of barley diseases [25]; and a support vector machine method to detect flowering rice in RGB images [11]. A support vector machine approach has also been used [27] to learn from SIFT features and a codebook generated over 7,500 images to classify higher plant taxa from images of leaves, achieving an accuracy of 72%.

Recently, a number of machine learning-focused approaches have taken part in the CVPPP challenge. One such approach used regression (specifically support vector regression) to count leaves in overhead views of rosette plants [10]. Leaf counting is again addressed in [17], which also uses machine learning to assist with segmentation via a Random Forest classifier. Other approaches have viewed the CVPPP leaf segmentation data set as an instance segmentation problem, segmenting each individual leaf in turn in the image. The most prominent paper in this area is one of the first applications of a recurrent neural network in this domain [20]. A spatial memory-equipped network allows the system to segment leaves one at a time and handle occlusion.

This brings us to the application of deep learning to plant phenotyping. Demonstrating the power of CNNs for classification, [3] develop a system, LeafNet, for taxon identification from images of leaves. This system outperforms previous approaches on standard image classification datasets. [18] use a CNN to classify a subset of plant features in small image sections. Localisation is performed by scanning each image and classifying overlapping sub windows. Consequently this approach is relatively slow, and lacks context due to the small cropped window sizes. Another approach has used a CNN-LSTM framework to classify plants into genotype[23]. The use of the LSTM is interesting, as it is used to improve classification over time; the author’s hypothesise is that growth rate is an important factor in determining genotype. The LSTM component does improve classification for the top-down rosette images used.

Of course, machine learning, and especially deep machine learning approaches are fuelled only by high quality, annotated datasets [24, 15]. For learning to be effective and efficient, the image data the computer is learning from must be both accurately captured and well annotated. This is our motivation for releasing our expertly-annotated dataset alongside the specific algorithms we have developed.

## 3. Method

In this section, we describe the new ACID dataset. We then discuss our network architecture that performs simultaneous feature localisation and classification, and our data augmentation and training approach.

### 3.1. The ACID Dataset

A key contribution of this work is a new dataset, containing images of wheat plants taken in glasshouse conditions. A doubled haploid population of spring wheat plants was obtained from the Nottingham/BBSRC Wheat Research Centre and grown in 2l pots in a glasshouse. This population was selected for its wide range in canopy and spike phenotypes. Imaging was conducted using a consumer grade 12MP camera fixed to custom-built imaging system providing a consistent black background. Each image contains multiple spikes at high resolution, with full annotation of the position of each spike, and further positions of each spikelet. Fig. 2 shows representative images from this dataset.

These images have been annotated by a single expert, at their native resolution of 1956×1530. The dataset is available at full resolution, with no augmentation or cropping. Each image is supplied with a JSON file containing coordinates of the base and tip of each ear (occasionally additional points should the ear be curved), and co-ordinates of all visible spikelets. Occluded spikelets were not annotated; however, partially occluded ears were left as continuous polylines rather than split up. This is a multi-instance dataset, in which each image contains multiple objects of the same class. In total there are 520 images, containing a total of 4,100 ears and 48,000 spikelets. Each image has also been tagged with the presence of an awned phenotype, (Fig. 2, right). Awned plants comprise about 1/3 of the dataset.

### 3.2. Network Architecture

Each image presented to the network could conceivably contain hundreds of similar objects to be localised and counted. Instead of predicting location directly, we perform pixel-wise regression, identifying areas of high-likelihood of each target. Detected features are then determined as the maximum points of this likelihood. This heatmap regression has been used successfully in, among other areas, human pose estimation [5]. Our network is based upon a stacked hourglass network [16], which itself is an evolution of fully-connected networks [9] and residual networks [13]. An outline of the network we use is shown in Fig. 3.

**Figure 3:**
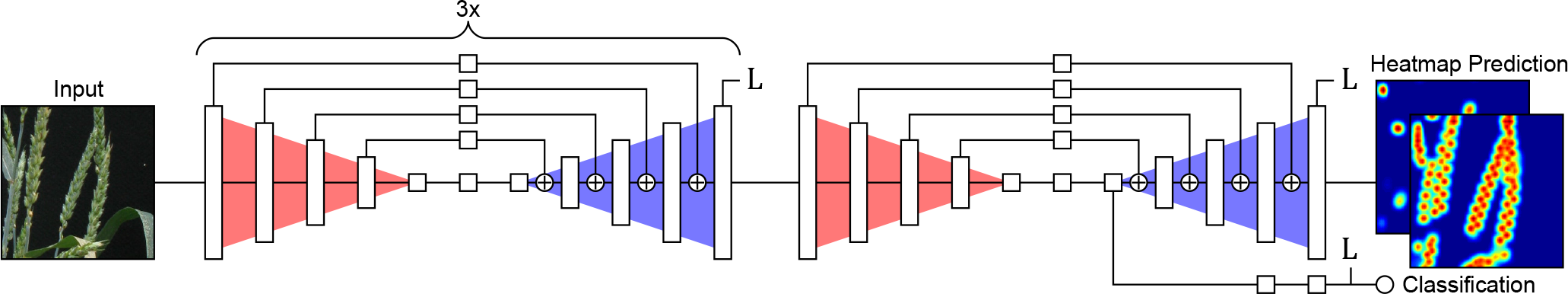
An overview of our CNN architecture containing 4 stacked hourglass networks.

The architecture of the network is based upon an encoding/decoding structure, in which a series of convolutional operations and spatial downsampling (red) begin by computing a fixed-size feature representation of the image. This feature space is then upsampled (blue) back to the original resolution, while lower-level features are re-combined in stages. Combining hierarchical features from multiple scales preserves spatial resolution in the network output.

Each white block in Fig. 3 represents one or more residual blocks, combining convolution and batch normalisation operations, and includes an additional skip layer that helps avoid vanishing gradients, aiding training in very deep networks such as this one. All residual blocks in the network output 256 features. The input to our network is an RGB image of size 256×256. A set of initial residual blocks (not shown in Fig. 3) reduces the spatial resolution to 64×64. The hourglass network operates at this lower resolution throughout. The output heatmaps are therefore also 64×64 pixels, with one for each feature being detected. We output two heatmaps in our network, trained on ear tips and spikelets separately. The network contains four stacked hourglasses, and includes intermediate supervision; the heatmap output at the end of each hourglass is used to calculate a loss, which guides training of the network.

The ACID dataset contains image-level labels specifying whether the imaged plant has an awned phenotype. We have extended our network to perform simultaneous classification by branching off the deep feature layer within the final hourglass. At this point, prior to upsampling, the data represents a spatially-invariant feature representation of the image. The branch contains two residual blocks, before a final convolutional block performing classification.

### 3.3. Data Augmentation

Our task is to locate and count wheat spikes and spikelets in the ACID dataset. Each image may contain a number of spikes, each of which will contain numerous spikelets. Both spikes and spikelets may appear very similar within a single image, but exhibit a large amount of variability between images, and lines. Due to hardware limitations, many deep network architectures have strict limits on image input size. This network has similar restrictions, where the input size cannot be increased far beyond 256px before GPU memory becomes the limiting factor. Many whole-image classification approaches will scale the image to the correct size during training or inference. In our case, each object of interest is small with respect to the original image, making global image scaling unwise. We wish to be able to accurately classify each large image completely, but preserve higher-resolution detail: each image must be split into regions.

Rather than splitting each image up prior to training, creating a fixed-size training set, we randomly crop images during training, with crops centred on spike positions. When an image is loaded, a random spike is chosen, and a training image produced at that location. The input to the network is 256×256; however, we have experimented with varying initial crop sizes, 256, 384, or 512 pixels. A larger initial crop that is eventually scaled to the correct input size will represent a wider field of view, with the features in the original image scaled down. This represents a compromise between the network seeing wider image context, and higher-resolution features. This does affect results, something we explore below. However, we use a default initial crop of 384 pixels, which represents a 2/3 scale of the original image resolution.

Additional random cropping, scaling, rotation and horizontal flipping is added to increase variability in the training set (Fig. 4). This means that when any given image is loaded, only a small part of that image is used during that training iteration. This means that a higher number of training epochs are required to capture the variability of the dataset, but that the eventual network should show increased performance.

Heatmap output is produced by performing identical transformations on the image labels, and then rendering each visible point as a two-dimensional Gaussian. The output heat maps are 64×64 pixels and we chose a standard deviation of 1.0 for spike tips, and 0.7 for the slightly smaller spikelets. In practice, we found any reasonable standard deviation was effective.

### 3.4. Training

The ACID dataset of 520 images was split into 415 training images, and 105 testing images. This split was performed at the image level, not the spike level, to ensure that no spikes from the same image could be seen in both training and testing sets. During training, random augmentation was applied as per Fig. 4 with random rotation and scaling drawn from normal distributions with standard deviations 0.25 radians and 25% respectively. Half of all input images were also horizontally flipped at random. No augmentation was performed on the testing image set. The network was trained end-to-end, from scratch, using RMSProp. We used a mean squared error loss function with an initial learning rate of 2.5×10^−4^, and reduced by a factor of 10 every 200 epochs. Training was run for 500 epochs, although performance usually plateaued around 300 epochs (Fig. 5).

**Figure 4:**
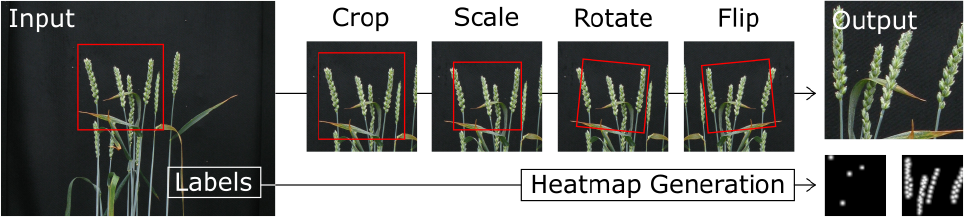
Data augmentation chooses an ear at random and creates a randomly transformed crop at that location. Any visible ground truth points are transformed in the same way.

**Figure 5:**
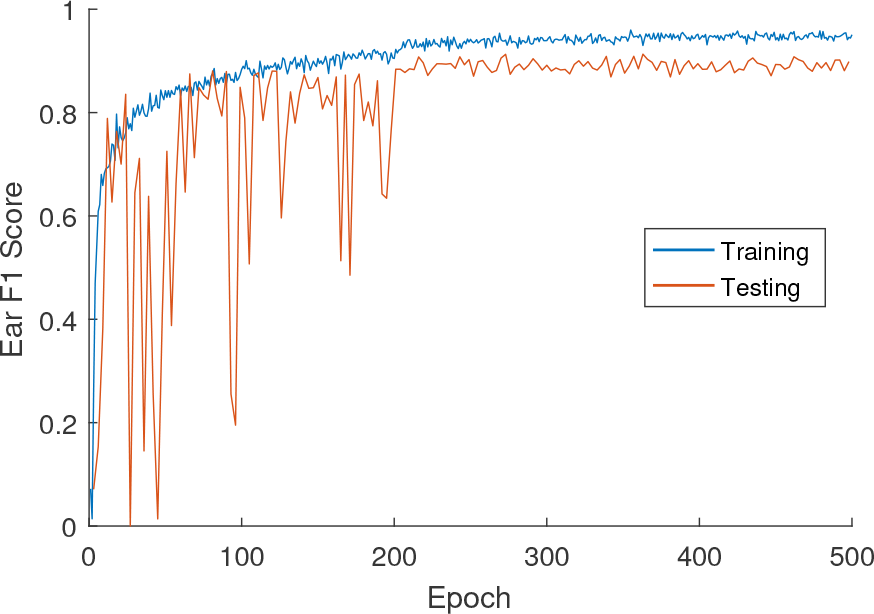
Ear accuracy over training and testing images throughout the training process.

Results below are presented on the final trained model after 500 epochs, not necessarily the best performing model during the entire training run. The time taken to complete each training run was approximately 3 hours. It is worth reiterating that due to our image augmentation mechanism, one epoch only represents a small view of the entire data, even though each image has been used once. This goes some way to explaining the large number of epochs required in this case.

The classification branch is trained in parallel using binary cross entropy. The classification branch loss is weighted at 5×10^−2^ compared to the heatmap MSE loss, to account for this BCE producing larger values in general, and so to avoid driving the training of the network entirely based on classification error.

### 3.5. Feature Localisation

Spike and spikelet positions are calculated from each output heatmap using non-maximal suppression (NMS). For all output pixels, any pixel with higher intensity than its four neighbours is classified as a feature. We have found that after sufficient training the network reliably positions local maxima at feature locations, with output distributions approximating the Gaussians used for training. The number of additional false positives generated by the NMS component of our approach is negligible.

## 4. Results

### 4.1. Evaluation

We evaluate our approach by calculating both the precision and recall for each network on the training set, along with the count accuracy for both spikes and spikelets. Precision represents the fraction of detections that are true positives. Recall represents the fraction of spikes and spikelets that have been correctly detected. It is common to combine these accuracy measures using the F1 score, as a general measure of performance.

What remains is to distinguish true positives from false positives, and determine which features have been correctly detected. We apply a distance threshold for successful detection, then vary this threshold to explore the efficacy of the approach at different tolerances. This normalised distance threshold is calculated relative to the size of the primary ear visible in each image (which is present in the annotations), see Fig. 8. A true positive for either a spike tip or spikelet is any predicted location that lies within this normalised distance of a ground truth point. Similarly, a false negative is any ground truth point that is not within the normalised distance of a predicted feature.

**Figure 8:**
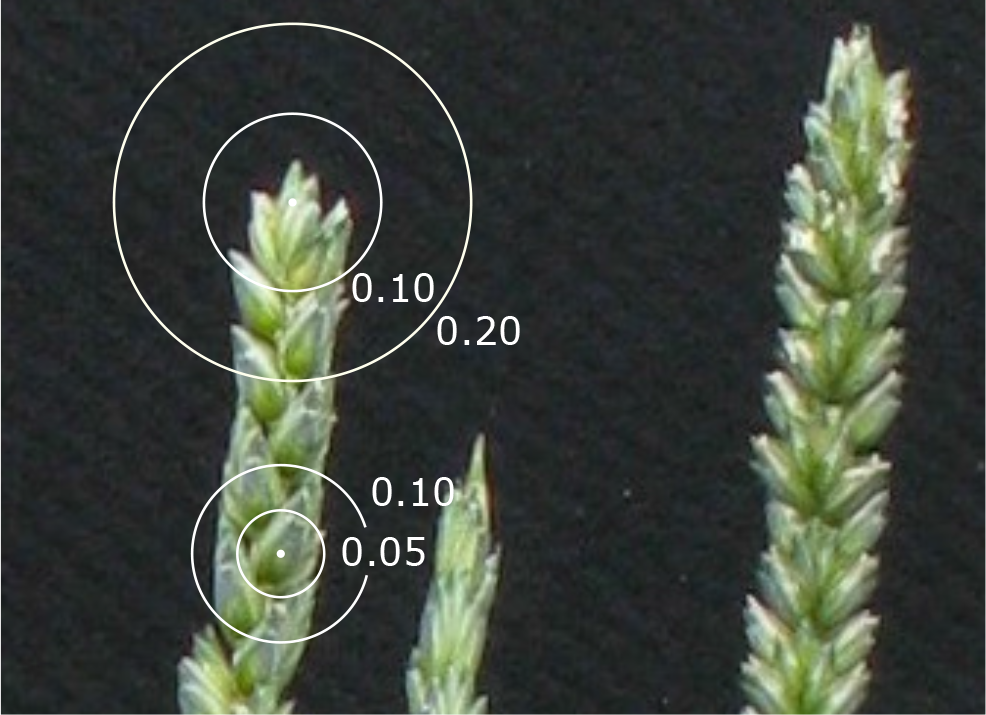
A visualisation of the normalised distances we used to measure accuracy. Here you can also clearly see individual spikelets within the spike

### 4.2. Numerical Results

Given the challenge of the problem, results achieved are very encouraging. Spikes are located with an F1 of 0.83@0.1 and 0.89@0.2, spikelets are located with an F1 of 0.88@0.05 and 0.96@0.1. For both features, we are confident that these normalised distances represent fairly strict tolerance for error. For comparison, an example wheat spike tip and spikelet marked with these normalised distances can be seen in Fig. 8. Variation in precision/recall over normalised distance can be seen in Fig. 7.

**Figure 7:**
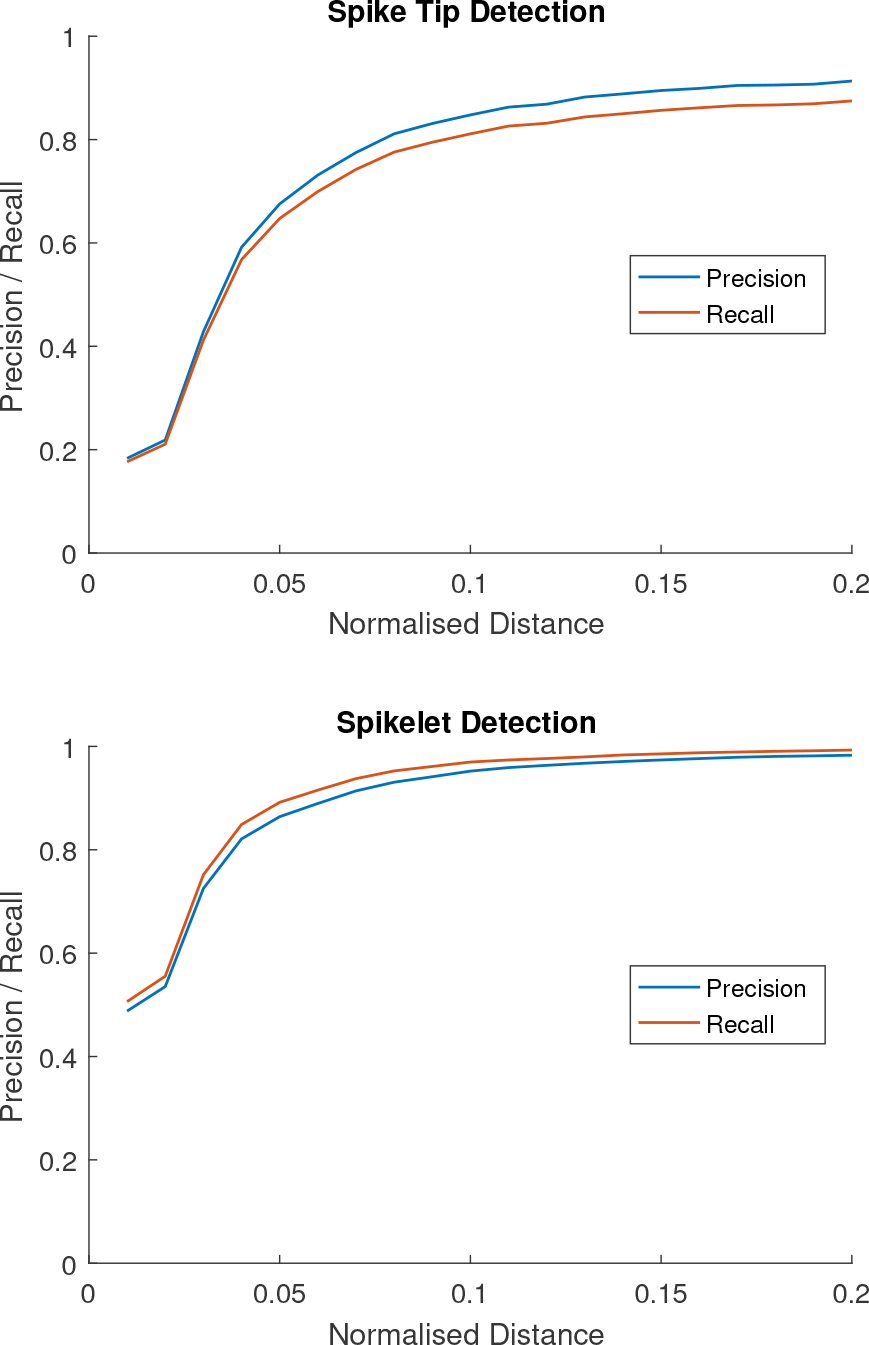
Precision and recall for the network as the normalised distance threshold is adapted.

#### Qualitative Analysis

Figures 1 and 9 show representative output from the network. The spike tip detection can be seen to work effectively in the presence of occlusion or large amounts of clutter. Spikelet detection is generally more accurate still, and is capable of distinguishing either dual rows of spikelets, or single rows when the spike is viewed rotated 90 degrees (see Fig. 9, lower right for examples of both). Detecting spike tips appears to be the harder problem. The failure modes on spike tips offer some insights into the network itself. While the network is quite capable of detecting tips even when occluded (Fig. 9), some occlusions will cause the detection to fail. We have also observed images in which the tip has still been detected, but has been incorrectly positioned at the edge of occlusion, rather than the true occluded location. This error will cause the recall and F1 scores to reduce, but the counting accuracy to remain the same.

**Figure 9:**
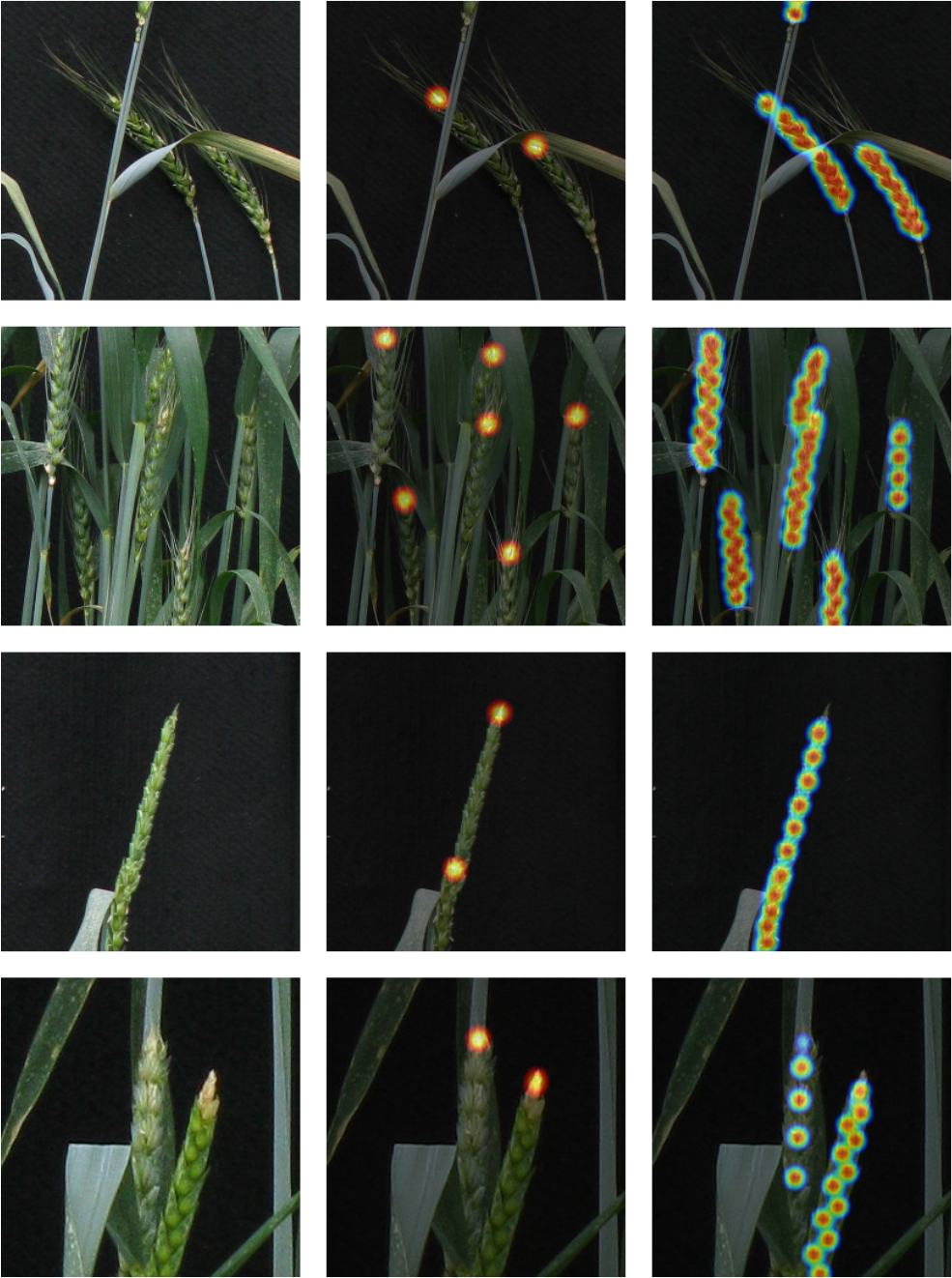
Some more challenging examples from the name dataset, including occlusion, background clutter, ambiguity and rotational asymmetry.

Where the tips of two spikes are very close together, the network will usually output two gaussian features very close together, or a single feature which is more spread out than usual on the heat map. In either case, this will adversely affect the counting accuracy, but will not affect our F1 measure, which is a weakness of our NMS feature extraction approach.

Spikelet detection may fail at the boundary between two overlapping spikes, due to the increased ambiguity between one and the other. This kind of overlap is not uncommon in the ACID dataset, but this also means multiple instances of this issue are included in the training set, offering the network some ability to distinguish between touching spikes.

#### Effect of Awned Phenotypes

We compared the testing F1 scores of all awned plants against all those that are not awned. We saw a marginal improvement of 1.3% F1 in testing accuracy for non-awned plants, suggesting awns are slightly more challenging to phenotype, however this was not a substantial difference, and may not be significant over many more images.

#### Effect of Augmentation

We trained the same network without data augmentation to measure the impact of a less varied training set. As expected, there was a degradation in performance for feature localisation on the testing set, particularly in spike tip detection. F1 reduced 7.5% for spikes, and 0.1% for spikelets. With the addition of random rotation and scaling to the testing set, this reduction increases to 8.1% for ears and 2.7% for spikelets. This suggests that in a non lab environment where image capture is less constrained, data augmentation will become even more important.

### 4.3. Input Image Resolution

Based on our own observation of the results, we believe that detection of wheat tips requires a great deal more context about the local image than individual spikelets. Intuitively, in a cluttered image containing multiple spikes, if a spike is partially cropped out of the input image, it will make accurate localisation of the tip harder. To confirm this hypothesis, we varied the input resolution to the network. As above, we initially crop a section of image for training or inference, perform augmentation, then if necessary the image is scaled to the size of the network input, 256×256. Altering the initial crop size is equivalent to scaling the entire image before it is used, and in essence changes the field of view available to the network.

We trained two additional networks, with input crop sizes of 256 and 512 pixels. The 256 crop input is native source resolution, where no image scaling is performed beyond that required by scale augmentation. Fig. 10 shows the results of this experiment, in which a smaller field of view performs notably worse on spike tip detection. In particular it is the recall ability of the network that is impaired, its ability to find tips, rather than too many false positives. Spikelet detection appears marginally better with a small field of view. This is perhaps also expected, spikelets are small features, and a 256pixel crop with no scaling preserves these at higher-resolution. However, it is also possible that the smaller crop size benefits from the normalised distance measure, with the size of an ear being larger with respect to the output heatmap when a smaller view is used. Nevertheless, the benefit is marginal, and has effectively disappeared when the normalised distance threshold reaches 0.1.

**Figure 10:**
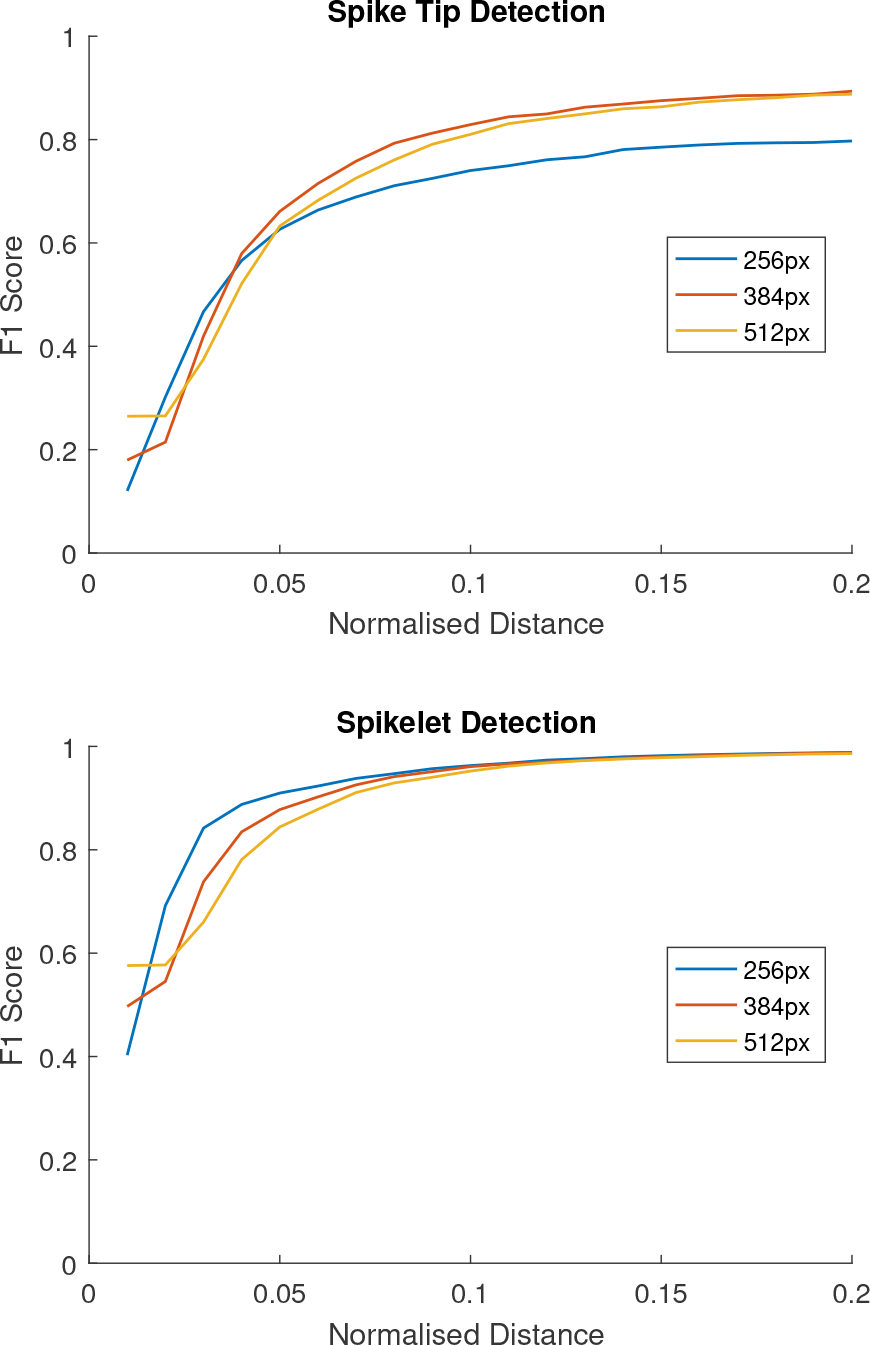
The effect of varying input resolution during data augmentation.

Given these results, it seems reasonable to recommend an input window size of 384 or 512 pixels. We measured the average length of spikes in the ACID dataset, and found this to be 235 pixels, with a large standard deviation of 68 pixels. Thus, either of these two resolutions offers a complete view of many spikes, improving tip localisation.

### 4.4. Counting Accuracy

To measure counting accuracy, we compared only the number of predicted features against the number of ground truth points and computed a percentage error, to simulate a count-based phenotyping task. These results were computed over 30 iterations of the testing set, to ensure that a representative sample of image crops was obtained. The results can be seen in Table 1.

**Table 1:** Percentage error for spike tip and spikelet counting at different input crop resolutions.

| Resolution (px) | Tip Error (%) | Spikelet Error (%) |
| --- | --- | --- |
| 256x256 | -14.46 | 0.0600 |
| 384x384 | -5.13 | 3.81 |
| 512x512 | -4.09 | 0.34 |

Given the lower recall performance on tips for the 256 pixel input size, the increased error is to be expected. It is interesting, however, that the 512 pixel input size performs well on both spike and spikelet counting, with very encouraging accuracy. This improvement over the smaller input sizes could be due to fewer spikes being truncated at the edges of the image, or a larger field of view adding context that helps resolve ambiguity.

The negative values for spike tip error indicate that the network tends to underestimate, rather than overestimate the number of tips. This follows from the results presented in Fig. 7, in which the precision of the network was higher than recall. Similarly, spikelet detection can slightly overestimate, where the recall of the network is higher than the precision

### 4.5. Classification

We have extended the network to perform simultaneous classification of awned plants. We frame the classification as the task of outputting a value close to one if the plant is awned, and close to zero if not. During inference, we threshold at 0.5 to convert the network branch output into a firm prediction. The classification accuracy for awns at the input resolutions we have examined are shown in Table 2. As above, results were computed over 30 iterations of the testing data.

Accuracy on heatmap generation of this network was unaffected by the addition to the architecture. As we might expect intuitively, classification accuracy increases as the window seen by the network increases. Nevertheless, all three resolutions offer extremely accurate classification of awned plants. Recognising awns is one of the easier classification challenges posed on wheat plants, compared with, for instance, growth stage classification. However, this acts as a demonstration that the hourglass design, if extended, can perform additional tasks beyond feature localisation. Our future work in this area will explore growth stage classification, flowering, and senescence.

**Table 2:**
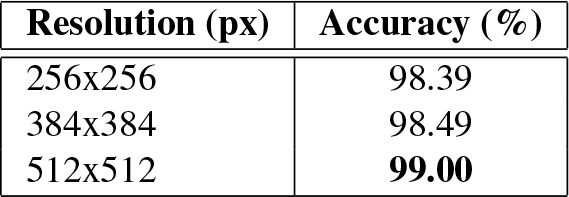
Awned classification accuracy at different input resolutions.

## 5. Conclusion

We have presented a new dataset, ACID, containing detailed annotations of wheat spikes and spikelets, as well as image level awn classification, on a varied phenotypic set of wheat lines. We have extended a deep convolutional neural network architecture to perform regression of feature locations, as well as image-level classification. We report very encouraging results on all aspects of the dataset. Our future work will focus on additional image-level classification, such as flowering plants, and retraining this network for field images, requiring additions to the dataset and application of transfer learning. Individual classification of spikelets (following promising work in [14]) will also be interesting to pursue, with a view to predicting yield. All data and code can be found at *http://plantimages.nottingham.ac.uk/*.

## References

[1] I. Arduini, L. Ercoli, M. Mariotti, and A. Masoni. Sowing date affect spikelet number and grain yield of durum wheat. Cereal Research Communications, 37(3):469–478, Sept. 2009.

[2] P. Armengaud, K. Zambaux, A. Hills, R. Sulpice, R. J. Pattison, M. R. Blatt, and A. Amtmann. Ez-rhizo: integrated software for the fast and accurate measurement of root system architecture. The Plant Journal, 57(5):945–956, 2009.

[3] P. Barr, B. C. Stver, K. F. Mller, and V. Steinhage. Leafnet: A computer vision system for automatic plant species identification. Ecological Informatics, 40:50–56, 2017.

[4] A. Blum. Photosynthesis and Transpiration in Leaves and Ears of Wheat and Barley Varieties. Journal of Experimental Botany, 36(3):432–440, Mar. 1985.

[5] A. Bulat and G. Tzimiropoulos. Human Pose Estimation via Convolutional Part Heatmap Regression, pages 717–732. Springer International Publishing, Cham, 2016.

[6] J. J. Casanova, S. A. O’Shaughnessy, S. R. Evett, and C. M. Rush. Development of a wireless computer vision instrument to detect biotic stress in wheat. Sensors, 14(9):17753–17769, 2014.

[7] F. Cointault, D. Guerin, J. Guillemin, and B. Chopinet. Infield triticum aestivum ear counting using colourtexture image analysis. New Zealand Journal of Crop and Horticultural Science, 36(2):117–130, 2008.

[8] A. French, S. Ubeda-Tomás, T. J. Holman, M. J. Bennett, and T. Pridmore. High-throughput quantification of root growth using a novel image-analysis tool. Plant Physiology, 150(4):1784–1795, 2009.

[9] R. Girshick, J. Donahue, T. Darrell, and J. Malik. Rich feature hierarchies for accurate object detection and semantic segmentation. In Proceedings of the IEEE conference on computer vision and pattern recognition, pages 580–587, 2014.

[10] M. V. Giuffrida, M. Minervini, and S. A. Tsaftaris. Learning to count leaves in rosette plants. Proceedings of the Computer Vision Problems in Plant Phenotyping (CVPPP), 2016.

[11] W. Guo, T. Fukatsu, and S. Ninomiya. Automated characterization of flowering dynamics in rice using field-acquired time-series rgb images. Plant Methods, 11(1):7, 2015.

[12] N. J. Halse and R. N. Weir. Effects of temperature on spikelet number of wheat. Australian Journal of Agricultural Research, 25(5):687–695, 1974.

[13] K. He, X. Zhang, S. Ren, and J. Sun. Deep residual learning for image recognition. In Proceedings of the IEEE Conference on Computer Vision and Pattern Recognition, pages 770–778, 2016.

[14] Y. Li, Z. Cui, Y. Ni, M. Zheng, D. Yang, M. Jin, J. Chen, Z. Wang, and Y. Yin. Plant density effect on grain number and weight of two winter wheat cultivars at different spikelet and grain positions. PLOS ONE, 11(5):1–15, 05 2016.

[15] G. Lobet. Image analysis in plant sciences: Publish then perish. Trends in Plant Science, 2017.

[16] A. Newell, K. Yang, and J. Deng. Stacked hourglass networks for human pose estimation. In European Conference on Computer Vision, pages 483–499. Springer, 2016.

[17] J.-M. Pape and C. Klukas. Utilizing machine learning approaches to improve the prediction of leaf counts and individual leaf segmentation of rosette plant images. In H. S. S. A. Tsaftaris and T. Pridmore, editors, Proceedings of the Computer Vision Problems in Plant Phenotyping (CVPPP), pages 3.1–3.12. BMVA Press, September 2015.

[18] M. P. Pound, A. J. Burgess, M. H. Wilson, J. A. Atkinson, M. Griffiths, A. S. Jackson, A. Bulat, G. Tzimiropoulos, D. M. Wells, E. H. Murchie, et al. Deep machine learning provides state-of-the-art performance in image-based plant phenotyping. bioRxiv, page 053033, 2016.

[19] G. J. Rebetzke, D. G. Bonnett, and M. P. Reynolds. Awns reduce grain number to increase grain size and harvestable yield in irrigated and rainfed spring wheat. Journal of Experimental Botany, 67(9):2573–2586, Apr. 2016.

[20] B. Romera-Paredes and P. H. S. Torr. Recurrent instance segmentation. CoRR, abs/1511.08250, 2015.

[21] P. Sadeghi-Tehran, K. Sabermanesh, N. Virlet, and M. J. Hawkesford. Automated method to determine two critical growth stages of wheat: Heading and flowering. Frontiers in Plant Science, 8:252, 2017.

[22] A. Singh, B. Ganapathysubramanian, A. K. Singh, and S. Sarkar. Machine learning for high-throughput stress phenotyping in plants. Trends in plant science, 21(2):110–124, 2016.

[23] S. Taghavi Namin, M. Esmaeilzadeh, M. Najafi, T. B. Brown, and J. O. Borevitz. Deep phenotyping: Deep learning for temporal phenotype/genotype classification. bioRxiv, 2017.

[24] S. A. Tsaftaris, M. Minervini, and H. Scharr. Machine learning for plant phenotyping needs image processing. Trends in plant science, 21(12):989–991, 2016.

[25] M. Wahabzada, A.-K. Mahlein, C. Bauckhage, U. Steiner, E.-C. Oerke, and K. Kersting. Metro maps of plant disease dynamicsautomated mining of differences using hyperspectral images. 10(1):e0116902.

[26] I. F. Wardlaw. Interaction between drought and chronic high temperature during kernel filling in wheat in a controlled environment. Annals of Botany, 90(4):469–476, Oct. 2002.

[27] P. Wilf, S. Zhang, S. Chikkerur, S. A. Little, S. L. Wing, and T. Serre. Computer vision cracks the leaf code. Proceedings of the National Academy of Sciences, 113(12):3305–3310, 2016.

